# FXR2 Promotes Autophagic Clearance of Sindbis Virus Capsid and Protects Against Viral Encephalitis

**DOI:** 10.64898/2026.10.23.761393

**Authors:** Vibha Singh, Phillip Dryden, Woo Yong Chang, Zhongju Zou, Yanghong Gu, David L. Nelson, Tao Wang, Josephine W. Thinwa, Tiffany A. Reese

## Abstract

Autophagy protects neurons from viral infection by removing toxic viral components, but the host factors that recognize viral proteins and direct their clearance remain incompletely defined. Here, we identify fragile X mental retardation syndrome-related protein 2 (FXR2) as a mediator of protective autophagy during Sindbis virus (SINV) infection. Quantitative proteomic analysis of host proteins associated with SINV capsid identified FXR2, an RNA-binding protein linked to stress granules and protein homeostasis. Co-immunoprecipitation and immunofluorescence confirmed that FXR2 associates and colocalizes with viral capsid in infected cells. Loss of FXR2 increased capsid accumulation, reduced LC3 lipidation and autophagosome formation, and impaired p62 degradation following infection. Reconstitution of FXR2-deficient cells restored autophagy and promoted capsid clearance. Although FXR2 deficiency did not significantly increase viral replication, it enhanced virus-induced cell death, suggesting that FXR2 protects infected cells by limiting viral protein toxicity rather than by directly restricting virus production. Consistent with this model, both Fxr2-null and heterozygous mice exhibited increased mortality following neurotropic SINV challenge, and Fxr2-null mice developed greater neuronal degeneration, inflammation, and cell death despite viral titers comparable to those in wild-type animals. Together, these findings identify FXR2 as a previously unrecognized component of the antiviral autophagy pathway that promotes viral capsid clearance and protects neural tissue from infection-associated damage.

## INTRODUCTION

Autophagy and stress granules are interconnected cellular stress responses that can strongly influence viral infection. Autophagy sequesters damaged organelles, protein aggregates, and invading pathogens in double-membrane vesicles for lysosomal degradation, thereby maintaining cellular homeostasis and, in some cases, restricting infection through the selective removal of viral components^1^. Stress granules are transient, membrane-free assemblies of untranslated messenger RNAs and RNA-binding proteins that form when translation initiation is inhibited and can concentrate antiviral sensors and signaling proteins. Autophagy also contributes to stress-granule clearance— a process termed granulophagy—helping prevent their persistent accumulation^2^. Viruses frequently manipulate both systems: some suppress autophagic degradation or stress-granule formation to evade host defenses, whereas others exploit autophagy-related membranes or stress-granule proteins to support viral replication^3,4^. Consequently, whether autophagy and stress granules promote antiviral defense or viral survival depends on the virus, host-cell type, stage of infection, and specific pathway components involved.

The targeting of viral proteins and particles for autophagosomal degradation is termed viral xenophagy, or virophagy^5^. Virophagy is an essential pathway for cell-intrinsic immunity against viruses that target nonrenewable cell populations, like neurons. For example, Sindbis virus (SINV), which is a positive-sense RNA alphavirus that causes viral encephalitis similar to other arthropod-borne viruses, is lethal in hosts that lack core autophagy machinery, such as Atg5^1^. One of the key features of autophagy-deficient SINV-infected neurons and brains, is accumulation of capsid protein and neuronal cell death^1,6^.

Fragile X–related (FMR) proteins—including FMRP, FXR1, and FXR2—and Ras-GAP SH3 domain-binding proteins 1 and 2 (G3BP1/2) are stress-granule-associated RNA-binding proteins that alphaviruses repurpose to support replication^7^. The viral nonstructural protein nsP3 binds these factors through motifs in its hypervariable domain, but the preferred partner differs among alphavirus lineages: New World encephalitic alphaviruses, such as Venezuelan equine encephalitis virus, interact directly with FMR-family proteins, whereas many Old-World alphaviruses, including SINV, chikungunya and Semliki Forest viruses, primarily recruit G3BP proteins^8^. These interactions redirect the proteins from canonical stress granules into nsP3-containing complexes, thereby suppressing aspects of the cellular stress response while concentrating viral RNA, replication machinery, ribosomes, and translation-initiation factors at sites of infection^9^. Recent evidence further suggests that nsP3 and these host RNA-binding proteins form biomolecular condensates that protect viral genomes from degradation and promote their translation and replication^10–12^. Thus, although FMR and G3BP proteins normally participate in cellular RNA regulation and stress responses, alphaviruses have evolved lineage-specific mechanisms to convert them into essential proviral factors.

Fragile X syndrome (FXS) is the most common heritable form of intellectual disability and autism in humans. Patients with FXS most commonly have an expansion of a poly-CGG trinucleotide repeat located in the 5′ UTR of the fragile X mental retardation 1 (FMR1) gene located on the X chromosome, resulting in hypermethylation and epigenetic silencing of the FMR1 gene^13^. It is well established that these mutations in FMR1 increase protein translation in neurons, and it was shown that mutant FMR1 neurons also have impaired autophagy^14^. Additional evidence indicates that the autosomal homologs to FMRP, FXR1 and FXR2, similarly participate in protein degradation pathways^15–18^.

In a proteomics screen aimed at isolating proteins that interact with SINV capsid, we identified FXR2, one of the stress granule-associated proteins described above. Interestingly, the FMR family of proteins were previously reported to not be essential for Old World alphavirus replication, whereas the G3BP stress granule proteins are required^8^. Therefore, we investigated whether FXR2 protects infected cells, thereby preventing tissue injury during SINV infection.

## RESULTS

### Identification of FXR2 in a proteomics screen for proteins that interact with Sindbis virus capsid

The degradation of viral components through virophagy plays a crucial role in viral protein clearance. However, the exact mechanisms for recognizing viral antigens and triggering autophagy remain poorly understood. To determine the time-point at which SINV infection initiates virus induced autophagy post-infection, we infected HeLa cells with SINV containing an HA-tagged capsid^6^ and examined autophagy induction over time. We observed HA-tagged SINV capsid as early as 1h post-SINV infection and increased accumulation 8h post-SINV infection (**Fig. 1A**). Further, we also probed for the autophagosome marker LC3-II conversion overtime post-SINV infection. We found that conversion of LC3-I to LC3-II increased overtime (0-8 h) along with the decrease in p62 protein, a key adaptor protein that recognizes ubiquitinated cargo including SINV capsid and links it to autophagosome membranes via LC3 binding (**Fig. 1A**)^1^. Based on these findings we selected post-SINV infection 1h as an early autophagic event and 7h as late autophagic event for further analysis SINV-induced autophagy.

**Figure 1.**
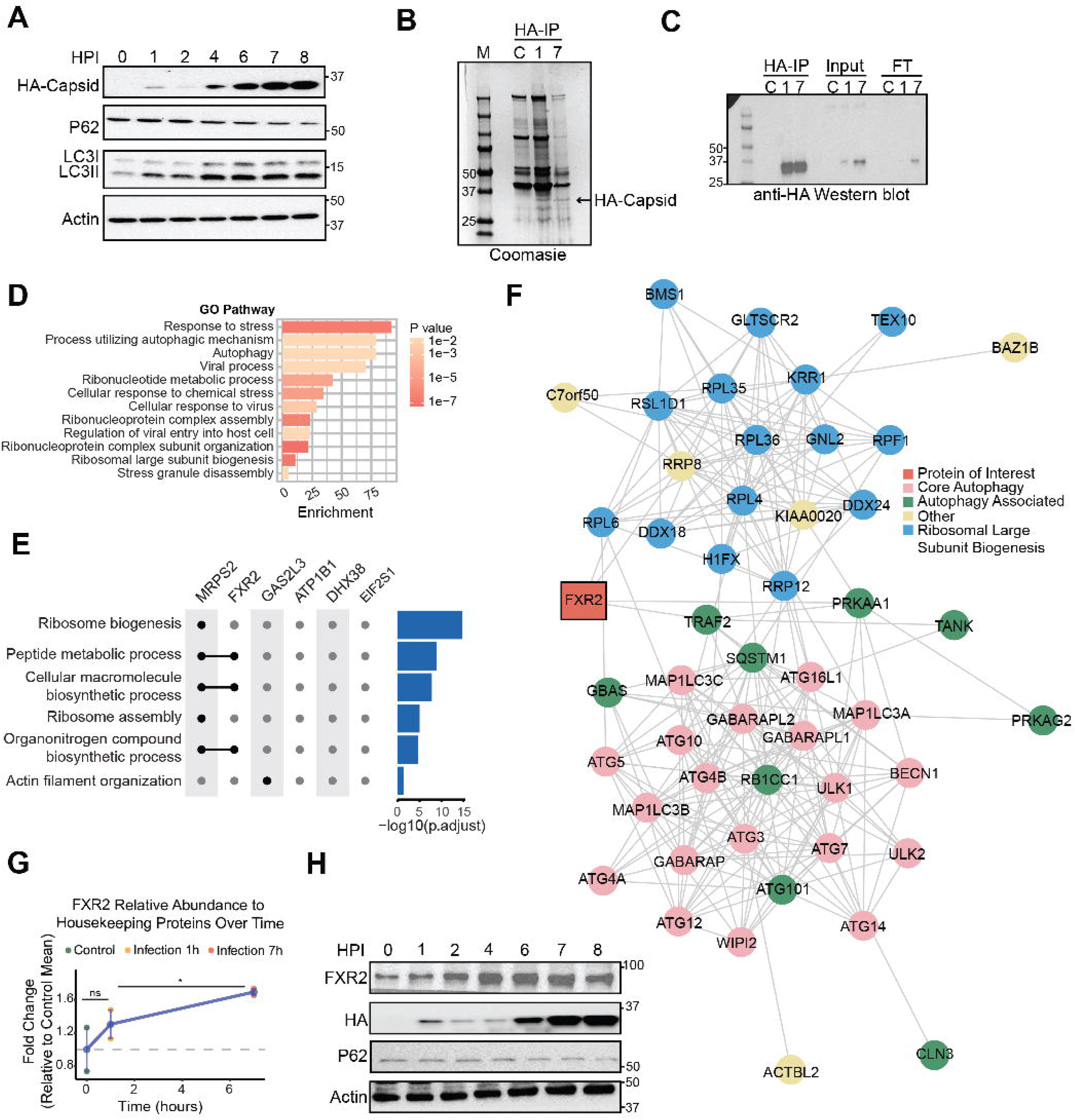
Identification of FXR2 as a Sindbis virus capsid-associated protein. HeLa cells were infected with SVIA.HA-capsid virus at MOI=10. **(A)** Representative immunoblot of HA-tagged capsid, p62, LC3-I/LC3-II, and actin in HeLa cells collected at the indicated hours post-infection (HPI). Actin served as the loading control. **(B)** HeLa cells were infected with SVIA.HA-capsid for 1 or 7 hours. Following lysis, control (C) or SVIA-infected lysates were subjected to HA immunoprecipitation. Coomassie-stained gel showing HA-capsid proteins in HA immunoprecipitation (HA-IP). **(C)** Immunoblot analysis using anti-HA antibody demonstrating enrichment of HA-capsid in HA-immunoprecipitated samples. HA-IP, input, and flow-through (FT) fractions collected from control (C) and 7-HPI samples are shown. **(D)** GO enrichment analysis of HA-capsid-associated proteins identified by mass spectrometry, highlighting stress, autophagy, viral response, ribonucleoprotein assembly, and ribosome biogenesis pathways. Bar length represents enrichment, and color represents the corresponding enrichment *p*-value. **(E)** Functional enrichment of selected HA-capsid-interacting proteins involved in ribosome biogenesis, peptide metabolism, macromolecule biosynthesis, and actin organization. The adjacent bars indicate statistical significance of pathway enrichment as -log10 (adjusted *P* value). **(F)** Protein-protein interaction network of the HA-capsid interactome. FXR2 is highlighted, with proteins grouped by autophagy, ribosome biogenesis, and other functional pathways. Connecting lines represent known or predicted protein-protein associations. **(G)** Relative FXR2 abundance by proteomic analysis following SINV infection, expressed as fold change relative to control. Values are expressed as fold change relative to the control mean. **(H)** Representative immunoblots showing changes in FXR2 abundance during the SINV infection time course. Cells were harvested at the indicated HPI and analyzed for FXR2, HA-capsid, p62, and actin. Immunoprecipitation followed by mass spectrometry was performed three times.

We applied an unbiased quantitative mass spectrometry-based proteomic approach to systematically identify host proteins that interact with the capsid protein of Sindbis virus. HeLa cells were infected with HA-tagged SINV capsid for 1h and 7h, followed by immunoprecipitation with anti-HA antibody. We confirmed abundance of HA-tagged SINV capsid in these samples both by Coomassie staining (**Fig. 1B, left panel)** and by western blot showing HA-tagged SINV capsid (**Fig. 1B, right panel).** These samples were subjected to LC-MS/MS for proteomics analysis.

By combining co-immunoprecipitation with high-resolution liquid chromatography–tandem mass spectrometry (LC-MS/MS), we mapped the host interactome of the SINV capsid proteins. For downstream functional analyses, we performed differential expression proteins and pathway enrichment using Gene Ontology Over Representation Analysis (GO-ORA) comparing control and SINV-Infected condition. Our findings revealed that the SINV capsid actively engages with a diverse set of host factors, many of which are components of key cellular pathways including *Response to stress*, *Cellular response to virus*, and *Viral process*. Of note, within the top 3 pathways with significant gene enrichment post-SINV infection, we found the pathways called *Process utilizing autophagic mechanism* and *Autophagy* (**Fig. 1D**) suggesting SINV-induced autophagic events.

To identify candidate proteins for functional validation, we compared our list of differentially interacting proteins to those identified a study that used siRNA knockdown and quantification of LC3 puncta to identify novel proteins required for autophagy ^19^. We identified six proteins MRPS2, FXR2, GAS2L3, ATP1B1, DHX38, and EIF2S1, that overlapped with the Orvedahl 2011 study. However, only FXR2, an RNA-binding protein called Fragile X Mental Retardation Syndrome-Related Protein 2, appeared in three significantly enriched GO pathways (adjusted p-value < 0.05): peptide metabolic process (adjusted p-value = 1.182e-09), cellular macromolecule biosynthetic process (adjusted p-value = 1.646e-08), and organonitrogen compound biosynthetic process (adjusted p-value= 2.628e-05) (**Fig. 1E**). In comparison, other candidate proteins showed much less significant pathway enrichment, with GAS2L3 appearing in only one pathway (actin filament organization), while ATP1B1, DHX38, and EIF2S1 showed no significant enrichment in any pathway. While MRPS2 showed presence in five enriched pathways, including ribosome biogenesis, ribosome assembly, and multiple metabolic processes, FXR2 was selected for further investigation due to its documented ability to selectively modulate mRNA translation in stress and autophagy contexts^20^.

We also employed Protein-Protein Interaction (PPI) network plot using STRING-analysis. Analysis of our PPI network revealed that FXR2, uniquely bridges two distinct functional modules: ribosomal large subunit biogenesis and core autophagy (**Fig. 1F**). This bridging function is mediated through interactions with proteins from both pathways, specifically TRAF2 and PRKAA1 (autophagy-related) and RRP8 and RSL1D1 (ribosome biogenesis-related). The topological position of FXR2 within our network suggests it may function as a molecular coordinator between these cellular processes (**Fig. 1F**).

Additionally, we examined FXR2 abundance over time in our proteomics data and found that FXR2 increased over time in response to infection (fold change = 1.62, one-sided p-value = 0.038) (**Fig. 1G**). The time course plot demonstrated a progressive increase in FXR2 protein expression following infection, with the most pronounced effect occurring at the later time point, suggesting a delayed but substantial response to infection. This finding was also validated by western blotting in HeLa cells infected with SINV from 0 to 8 h.p.i. showing an increase in FXR2 protein expression (**Fig. 1H**).

### FXR2 associates with SINV capsid

FXR2 is predominantly localized in the cytoplasm, where it associates with polyribosomes and messenger ribonucleoprotein (mRNP) complexes^21^. It can also shuttle into the nucleus and participate in RNA processing, although it’s less abundant in the nucleus compared to the cytoplasm^22,23^. This dynamic localization supports FXR2’s role in RNA binding, translation regulation, and cellular stress responses—functions that may be hijacked during viral infections such as SINV to facilitate processes like virophagy. To examine the subcellular distribution of FXR2 following SINV infection (MOI 10, 8 hour), we performed nuclear and cytoplasmic fractionation using a commercially available kit. Western blot analysis of these fractions revealed that the majority of FXR2 localized to the cytoplasmic fraction compared to the nuclear fraction (**Fig. 2A**) post-SINV infection. Lamin B1 and vinculin were used as markers for nuclear and cytoplasmic fractionation, respectively. In addition to fractionation, we employed indirect immunofluorescence (IF) to visualize FXR2 distribution at 6 hours post-SINV infection.

**Figure 2.**
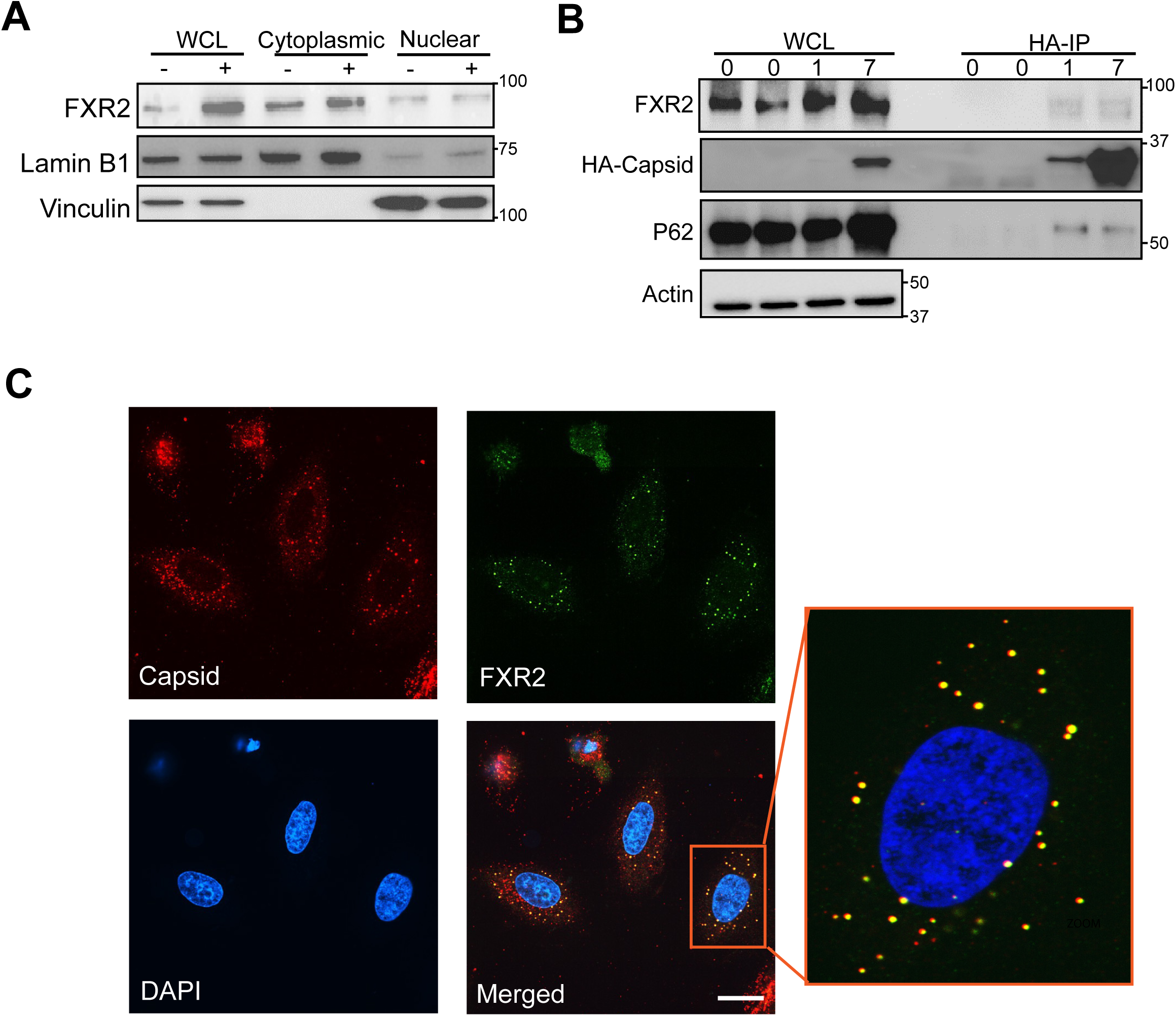
FXR2 associates with SINV capsid-containing protein complexes and localizes with viral capsid during infection. **(A)** Representative immunoblot analysis of whole-cell lysates (WCL) and HA-immunoprecipitated (HA-IP) fractions from cells collected at 0, 1, and 7 HPI. Samples were analyzed for FXR2, HA-capsid, and p62. Actin in the WCL served as a loading control. Enrichment of FXR2 and p62 in HA-capsid immunoprecipitates was assessed with SINV infection overtime. **(B)** Subcellular fractionation analysis of FXR2 in control (-no infection) and SINV-infected (+) cells. FXR2 abundance was examined in whole-cell lysate, cytoplasmic, and nuclear fractions. Lamin B1 and vinculin were used as nuclear and cytoplasmic fraction markers, respectively. **(C)** Representative immunofluorescence images showing the cytoplasmic localization of SINV capsid (red) and FXR2 (green) in infected cells. Nuclei were counterstained with DAPI (blue). The merged image and enlarged region demonstrate areas in which capsid and FXR2 signals are present in close proximity. Bar represents 20 μm. Each panel represents 2 independent experiments.

We next confirmed by immunoprecipitation that FXR2 associates with SINV capsid. Cells were infected with SINV-HA, and capsid was immunoprecipitated at 1 and 7 hours after infection. We detected both FXR2 and p62 in samples after HA-capsid immunoprecipitation (**Fig. 2B**). Immunofluorescence analysis confirmed cytoplasmic enrichment of FXR2 in infected cells (**Fig. 2C, zoomed inset).** Further, when we examined colocalization of capsid and FXR2 by immunofluorescence, we found that capsid colocalizes with FXR2 (**Fig. 2C**). These data corroborate our proteomics analysis that indicated that FXR2 associates with capsid.

### Loss of FXR2 leads to increased sindbis virus capsid accumulation during virus infection

Because SINV capsid protein accumulates in the absence of autophagy ^1,6^, we first examined if SINV capsid protein accumulated more in cells deficient in FXR2. Using an siRNA approach, we knocked down FXR2 (**Fig. 3A**). HeLa cells transfected with either control siRNA or siRNA to FXR2 were infected with SINV expressing an mCherry-tagged capsid protein^1^. In FXR2 knockdown cells, we observed more cells with mCherry capsid compared to control siRNA expressing cells (**Fig. 3B, C**). We subsequently made HeLa knockout clones that were deficient in FXR2, using CRISPR/Cas9 technology (**Fig. 3D**). Similar to experiments with siRNA knockdown, FXR2-KO cells accumulated more mCherry-capsid compared to wildtype cells (**Fig. 3E, F**). These data suggest that loss of FXR2 impairs viral capsid clearance, leading to capsid accumulation in infected cells.

**Figure 3.**
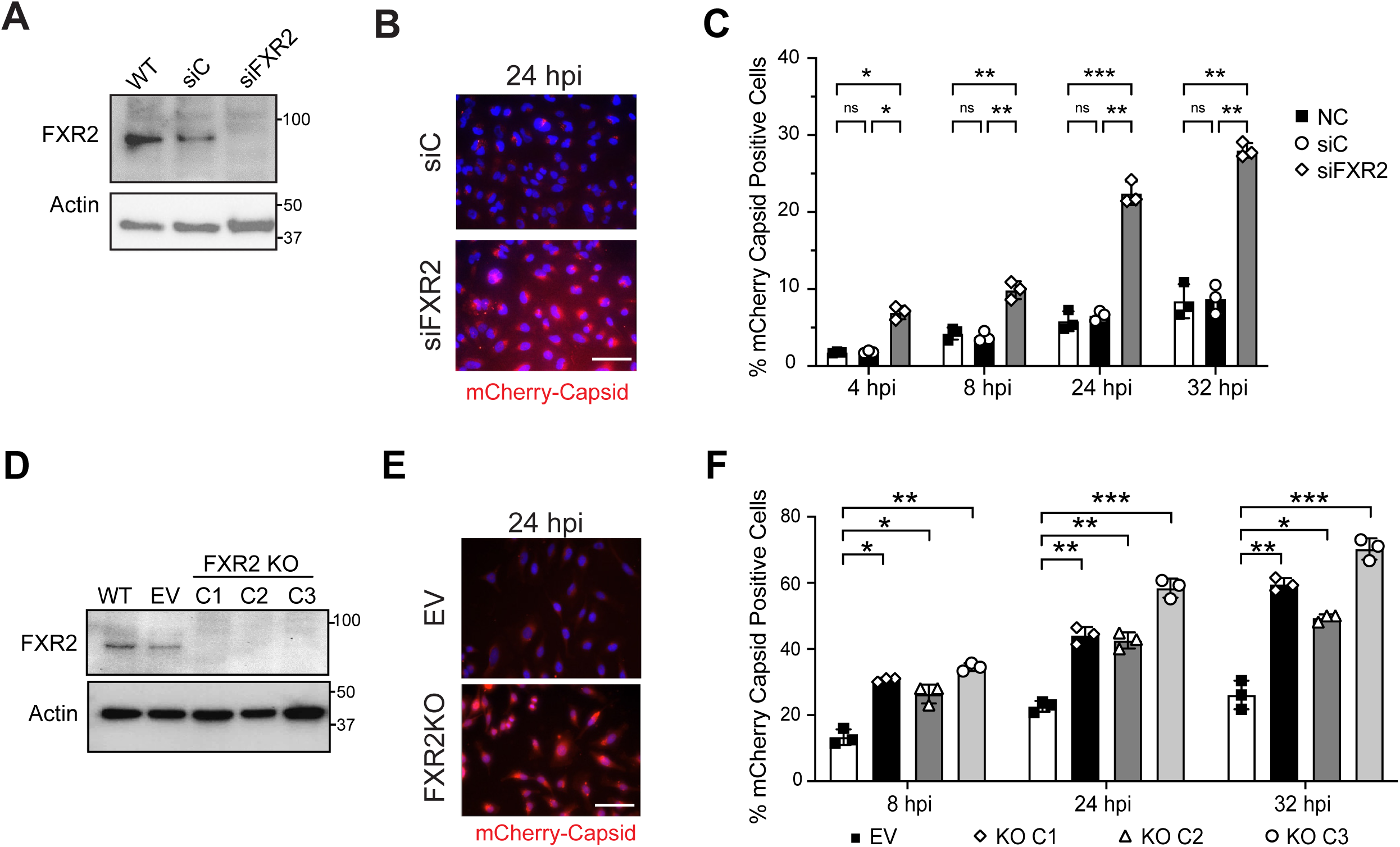
Depletion of FXR2 induces SINV capsid accumulation. **(A)** Representative immunoblot confirming siRNA mediated depletion of FXR2. FXR2 protein expression was compared among control siRNA (siC) and FXR2-targeting siRNA (siFXR2) conditions, with actin used as a loading control. **(B)** Representative fluorescence microscopy images of cells transfected with SiC or siFXR2 and infected with SVIA.mCherry-capsid. Images were acquired at 24 HPI. Viral capsid is shown in red and nuclei are shown in blue. Scale bar: 100 μm **(C)** Quantification of mCherry-capsid-positive cells at 4, 8, 24, and 32 HPI following treatment with non-targeting control (NC), control siRNA (SiC), or FXR2-targeting siRNA (siFXR2). **(D)** Representative immunoblot confirming FXR2 loss in independently generated FXR2-knockout (KO) cell clones (C1-C3) compared with wild-type (WT) and empty-vector (EV) controls. Actin served as the loading control. **(E)** Representative fluorescence microscopy images of empty-vector control and FXR2-KO cells infected with SVIA.mCherry-capsid and imaged at 24 HPI. Viral capsid is shown in red and nuclei in blue. Scale bar: 100 μm. **(F)** Quantification of mCherry-capsid-positive cells in EV control cells and three independently generated FXR2-KO clones at 8, 24, and 32 HPI. Individual symbols represent three experimental replicates. Statistical differences were assessed using a two-way ANOVA with a mixed-effects model followed by Tukey’s multiple-comparisons test. Statistical significance is indicated by asterisks: *\*p* <0.05, **p<0.01, ***p <0.001, and ****p <0.0001.

### Loss of FXR2 impairs autophagy induction

In addition to capsid accumulation, cells deficient in autophagy have impaired conversion of LC3-I to the autophagosome-associated, LC3-II, and accumulation of p62^24^. HeLa cells were transfected with siRNAs for FXR2 and infected with SINV-HA-capsid virus. In **Figure 4A**, we confirmed knockdown of FXR2 from 0 to 8 hours post SINV infection. We examined the accumulation of capsid in control and knockdown cells and observed increased capsid accumulation in FXR2 knockdown cells at 6 and 8 hours post infection, in agreement with Figure 3. We monitored p62 levels and saw that cells transfected with control siRNA had decreased p62 over the timecourse of infection, but that FXR2 knockdown cells maintained a higher level of p62, suggesting a defect in virus-induced autophagy in FXR2 knockdown cells (**Fig. 4A**).

**Figure 4.**
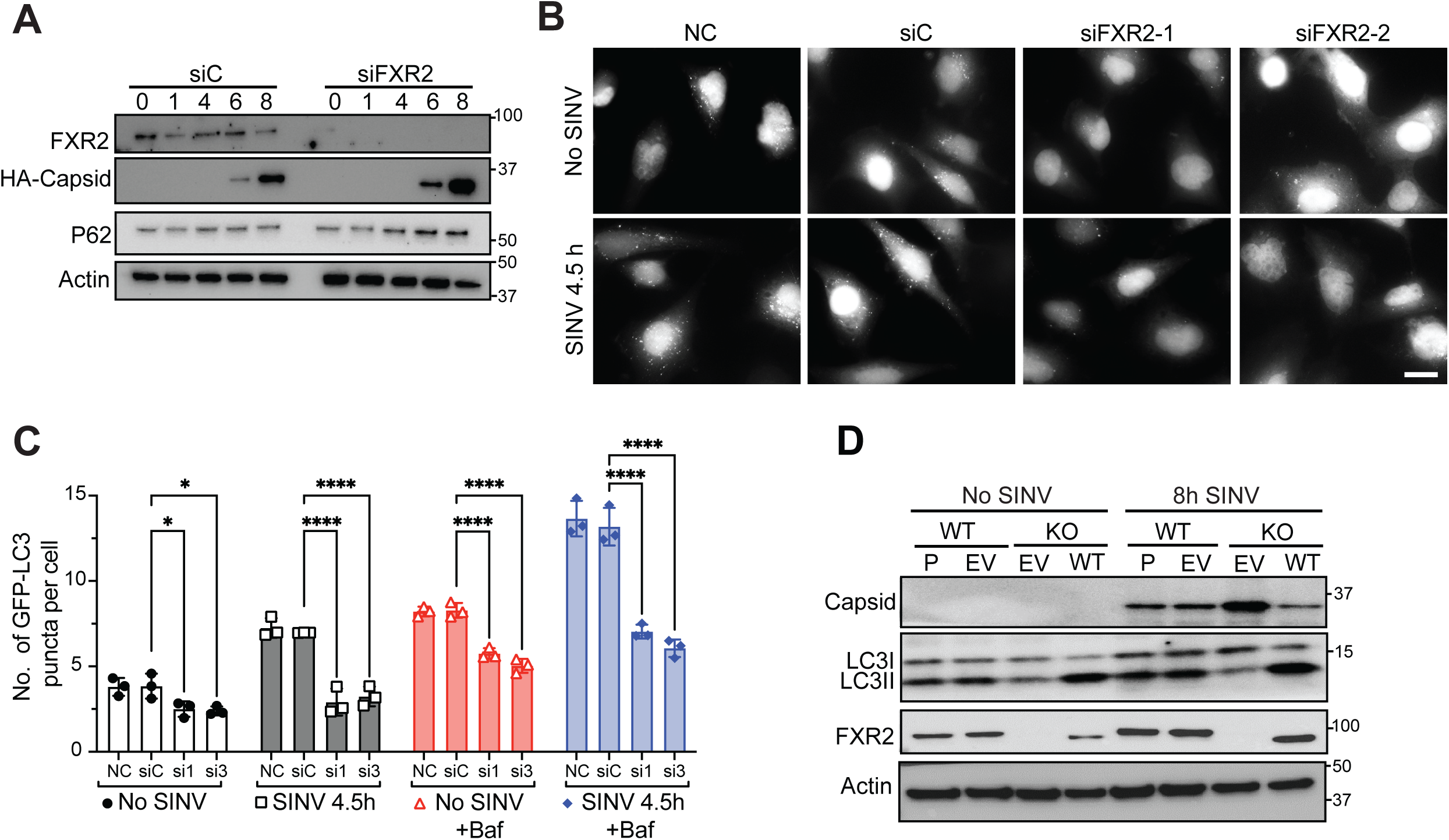
Loss of FXR2 impairs autophagy induced by SINV infection. **(A)** Representative immunoblot analysis of FXR2, HA-capsid, p62, and actin in cells treated with control siRNA (siC) or siFXR2 and collected at 0, 1, 4, 6, and 8 HPI. Actin served as the loading control. Represents 3 independent experiments. **(B)** Representative fluorescence microscopy images of GFP-LC3 puncta in HeLa cells treated with non-targeting control (NC), control siRNA (siC), or two independent FXR2-targeting siRNAs (siFXR2-1 and siFXR2-2). GFP-LC3 puncta images were captured under no infection and at 4.5 h following SINV infection. Scale bar: 20 μm **(C)** Quantification of GFP-LC3 puncta per cell in control and FXR2-depleted cells under no infection or following 4.5 h of SINV infection, in the absence or presence of bafilomycin A1 (Baf). Individual symbols represent three different replicates. Statistical differences were assessed using a two-way ANOVA with a mixed-effects model followed by Tukey’s multiple-comparisons test. Statistical significance is indicated by asterisks: *P* <0.05, \**P*<0.01, \*\**P* <0.001, and \*\*\**P* <0.0001. **(D)** Representative immunoblot examining LC3 processing following SINV infection in wild-type (WT) and FXR2-knockout (KO) cells. FXR2-KO cells were reconstituted with WT FXR2 or the corresponding control vector. Cells were analyzed with no infection or at 8 HPI for capsid, LC3-I/LC3-II, FXR2, and actin. Represents 3 independent experiments.

One hallmark feature of autophagy is the formation of LC3 puncta in the cytoplasm of cells. To measure the formation of LC3 puncta^24^, HeLa cells stably expressing GFP-LC3 were transfected with control or FXR2 siRNA knockdown constructs and infected with SINV. SINV infection induced GFP-LC3 puncta 4.5 hours post infection (**Fig. 4B, C**). In contrast, the FXR2 siRNA expressing cells had decreased puncta formation prior to infection, and more pronounced defect in LC3 puncta formation 4.5 hours post SINV. Of note, treatment of cells with bafilomycin A1 (BafA1), a lysosomal acidification and autophagic flux inhibitor^24^, led to increased LC3 puncta at baseline and after SINV infection in both control and FXR2 knockdown cells (**Fig. 4C**).

We examined whether FXR2-KO cells accumulated capsid and had defective conversion of LC3-I to LC3-II, and whether add-back of FXR2 could rescue the phenotype. In agreement with siRNA knockdown of FXR2, FXR2-KO cells accumulated more capsid after SINV infection compared to parental (P) HeLa cells (**Fig. 4D**). FXR2-KO cells (KO-EV) also had impaired conversion of LC3-I to LC3-II 8 hours post infection, in contrast to wildtype (WT-P) HeLa cells. Notably, reconstitution of FXR2-KO cells with WT FXR2, compared to empty vector (EV)-treated cells, restored conversion of LC3-II and promote degradation of capsid (**Fig. 4D**). Together, these data support that FXR2-deficient cells infected with SINV have altered autophagy.

### Loss of FXR2 increases cell death after SINV infection

Because we observed increased viral capsid protein, we next tested if FXR2 knockout cells had increased viral replication. Wildtype and FXR2-KO cells were infected with MOI=0.01 SINV and viral growth was quantified by plaque assay over time. We observed only a modest, but not statistically significant increase in virus replication in knockout cells compared to wildtype cells, suggesting that FXR2 deficiency does not promote increased virus replication (**Fig. 5A**). This agrees with previously published work showing that loss of individual or a combination of FMR family of proteins does not alter SINV replication^8^.

**Figure 5.**
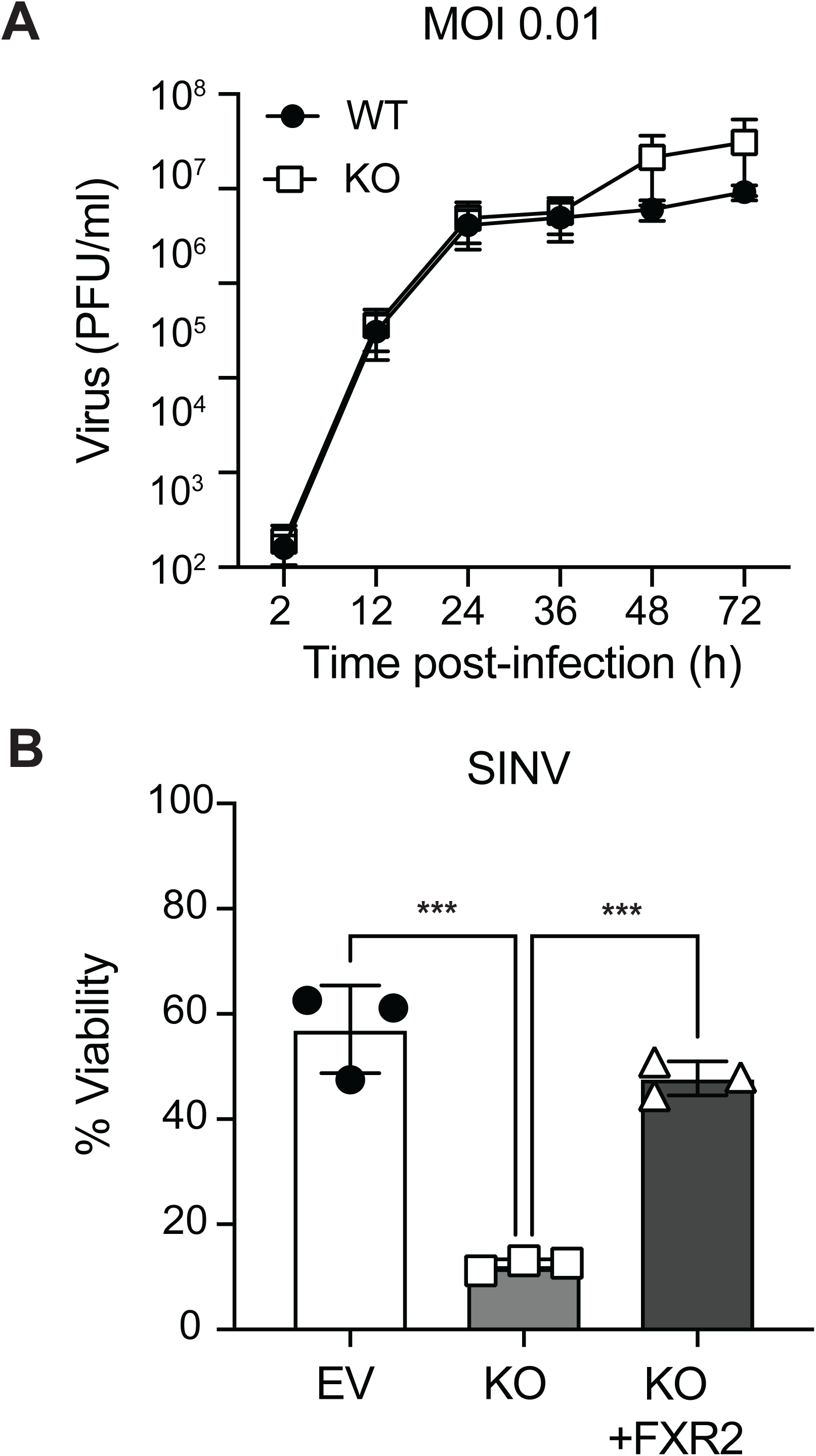
FXR2 deficiency alters SINV infection-associated cytotoxicity without altering viral production. **(A)** Multi-step SINV growth kinetics in wild-type (WT) and FXR2-knockout (KO) HeLa cells infected at an MOI of 0.01. Culture supernatants were collected at the indicated times post-infection, and infectious virus production was determined by plaque assay and expressed as plaque-forming units per milliliter (PFU/ml). Represents 2 independent experiments. **(B)** Cell viability following SINV infection in empty-vector control (EV), FXR2-KO, and FXR2-KO HeLa cells reconstituted with FXR2 (KO+FXR2). Viability is expressed as a percentage of the corresponding control. Individual symbols represent three independent experiments.

Accumulation of viral proteins and defective autophagic clearance of these proteins is associated with cell toxicity^1^. To determine if loss of FXR2 impacted cell survival, WT and FXR2-KO HeLa cells were infected with SINV and cell viability was measured 24 hours after infection (**Fig. 5B**). Loss of FXR2 significantly decreased cell viability after SINV infection, and complementation of FXR2 restored viability to levels comparable to wildtype infected cells. These data suggest that FXR2 promotes cell survival after SINV infection.

### FXR2 is necessary for survival of mice after sindbis virus challenge

Because autophagy is required for host survival after infection with multiple neurotropic viruses^1,6^, we next asked if FXR2 was required for survival after SINV challenge. Because the original Fxr2-knockout mouse maintained a neomycin cassette and was not a complete null for FXR2 protein^25^, we used a newly developed *Fxr2*-knockout mouse that does not have a neomycin insertion and is a complete null (**Fig. 6A**). Neonatal *Fxr2^-/-^*, *Fxr2^+/-^*, and *Fxr2^+/+^*mice were challenged with SINV intracranially. Compared with wildtype mice, *Fxr2^-/-^* mice were markedly more susceptible to SINV challenge (**Fig. 6B**). Interestingly, *Fxr2^+/-^*heterozygous littermates were also more susceptible to SINV challenge, with an intermediate phenotype between knockout and wildtype, indicating that two functional copies of *Fxr2* are required for efficient control of SINV infection. Although there was a notable difference in survival between wildtype and deficient mice, when we collected brain tissue and examined virus replication by plaque assay, we found no significant difference in viral titer in wildtype and knockout mice (**Fig. 6C**). This observation agrees with our *in vitro* replication data (**Fig. 5A**), suggesting that FXR2 does not play a role in limiting virus replication, but may instead be important for maintaining tissue homeostasis.

**Figure 6.**
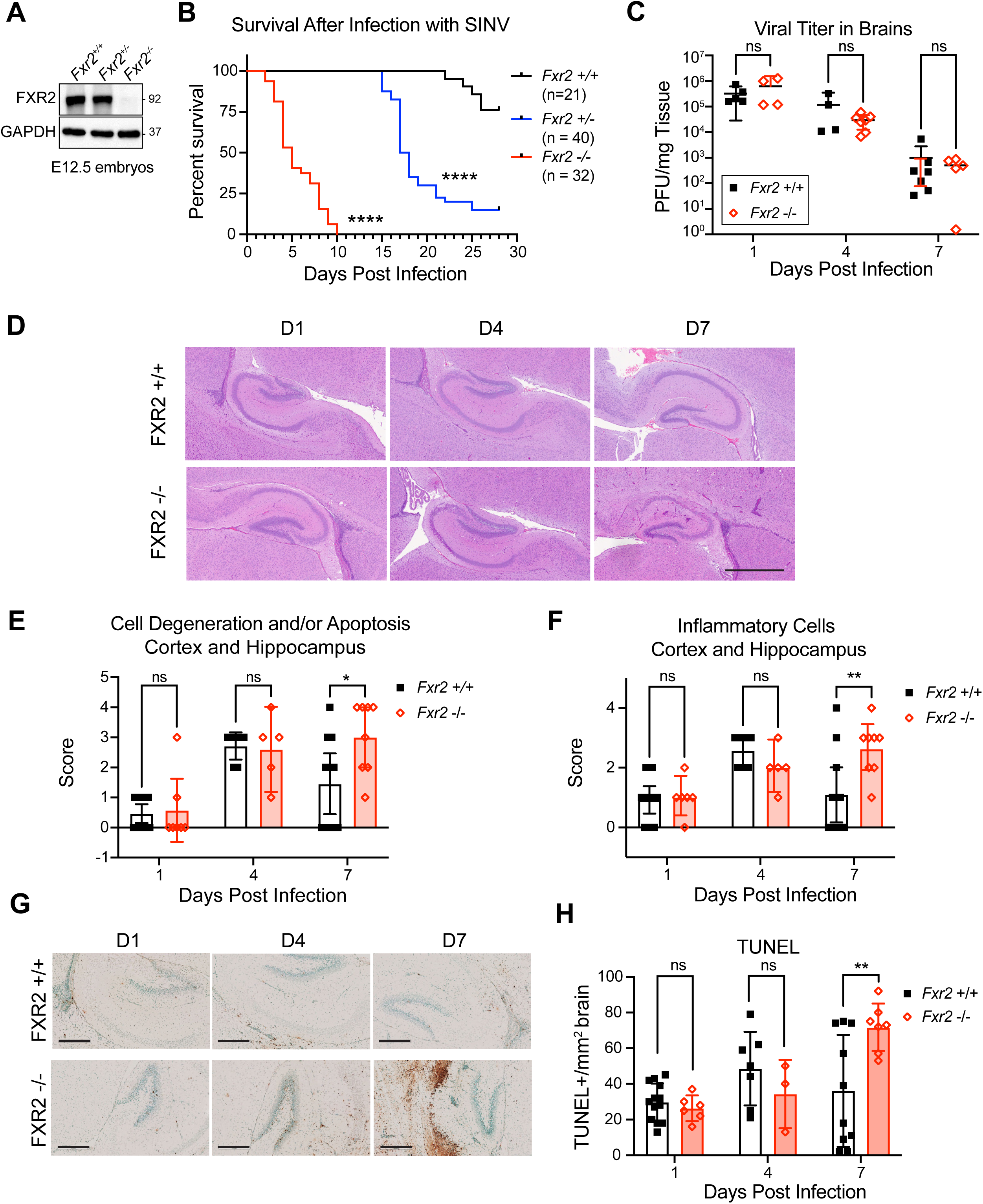
Loss of FXR2 increases susceptibility and brain pathology after SINV infection. **(A)** E12.5 embyos from *Fxr2^-/-^*, *Fxr2^+/-^*, and *Fxr2^+/+^* were analyzed for FXR2 protein expression. **(B – H)** Seven-day-old *Fxr2^-/-^*, *Fxr2^+/-^*, and *Fxr2^+/+^*littermates were infected with 1 × 10³ PFU of the dsTE12Q strain of SINV. **(B)** Kaplan-Meier survival analysis of FXR2 wild-type (*Fxr2+/+*, *n*=21), heterozygous (*Fxr2+/-*, *n*=40), and knockout (*Fxr2-/-, n*=32) mice following SINV infection. Survival was monitored for up to 30 days post-infection. Significance was assessed using Log-rank (Mantel-Cox) test using Prism. **** p<0.0001. **(C)** Viral burden in brains of wildtype *Fxr2^+/+^*and knockout *Fxr2^-/-^* mice at days 1, 4, and 7 post-infection. Viral titers were determined by plaque assay and expressed as plaque forming units (PFU) per mg of brain tissue. Individual symbols represent individual animals. ns, not significant. **(D)** Representative hematoxylin and eosin (H&E) stained brain sections from wildtype *Fxr2^+/+^* and knockout *Fxr2^-/-^* mice collected at days 1, 4, and 7 post-infection. **(E)** Histopathological scoring of neuronal/cellular degeneration and/or apoptosis in the cortex and hippocampus of wildtype *Fxr2^+/+^*and knockout *Fxr2^-/-^* mice at days 1, 4, and 7 following infection. **(F)** Histopathological scoring of inflammatory-cell infiltration in the cortex and hippocampus of wildtype *Fxr2^+/+^* and knockout *Fxr2^-/-^* mice at the indicated days post-infection. **(G)** Representative TUNEL-stained brain sections from wildtype *Fxr2^+/+^*and knockout *Fxr2^-/-^* mice at days 1, 4, and 7 after SINV infection. TUNEL-positive cells are visualized by brown staining. **(H)** Quantification of TUNEL-positive cells per mm² of brain tissue in wildtype *Fxr2^+/+^*and knockout *Fxr2^-/-^* mice at days 1, 4, and 7 post-infection. Individual symbols represent individual animals. Results for C-G were analyzed using Mixed-effects analysis on Prism with Tukey’s multiple comparison test. * p<0.05, ** p<0.01, **** p<0.000, n.s. not significant.

To further examine the tissue after SINV infection, brain tissue was collected and examined by histology. Histopathological analysis and scoring of H&E-stained tissue revealed an increase in cell degeneration and/or apoptosis in the cortex and hippocampus 7 days after infection in knockout animals (**Fig. 6D, E**). Furthermore, *Fxr2^-/-^*hippocampus and cortex had increased inflammation 7 days after SINV infection **(Fig. D, F)**.

To directly measure cell death, we stained tissue with TUNEL. Quantification of TUNEL staining revealed increased cell death and tissue destruction of the brains of *Fxr2^-/-^* compared with wildtype mice at day 7 post infection (**Fig. 6G, H**). Taken together, these data indicate that FXR2 is an essential gene that prevents cell death and tissue destruction after SINV infection of the brain.

## DISCUSSION

Here we report an unexpected role for FXR2 as a host factor that limits cellular injury during Sindbis virus infection. FXR2 associated with viral capsid in infected cells, and FXR2-deficient cells accumulated more capsid, showed altered autophagy markers, and had reduced viability. In mice, loss of Fxr2 increased brain injury and mortality without a significant increase in the viral titers measured. Together, these findings suggest that FXR2 helps preserve tissue integrity during infection. The cell experiments implicate autophagy and capsid clearance in this protection, although whether the same pathway accounts for the phenotype in infected neurons remains to be determined.

The increased mortality and neuropathology of *Fxr2*-deficient mice occurred without a corresponding increase in brain viral titers, indicating that FXR2 primarily promotes host tolerance to infection rather than resistance to viral replication. This phenotype resembles the protective effects of autophagy previously observed during SINV encephalitis, in which defective clearance of viral capsid increases neuronal injury and mortality without necessarily increasing viral burden ^1,6^. Because neurons are long-lived and have limited regenerative capacity, efficient disposal of abundant or aggregation-prone viral proteins may be particularly important for preserving tissue function during infection. The intermediate susceptibility of Fxr2-heterozygous mice further suggests that this protective activity is sensitive to FXR2 dosage. Future studies using neuron-specific deletion of Fxr2 and analyses of inflammatory and cell-death pathways should clarify whether FXR2 acts principally within infected neurons or also modifies responses in microglia, astrocytes, or infiltrating immune cells.

Previous reports indicate that alphaviruses hijack stress granule proteins, such as the FMR and G3BP proteins, and co-opt them for viral replication. Interestingly, the Old-World alphaviruses, including SVIA and chikungunya virus, require G3BP proteins to support virus replication, but the FMRP proteins are not required. When all three FMRP proteins (FMRP1, FXR1, and FXR2) are knocked out in cell lines, virus replication is equivalent to wildtype, similar to our results in single FXR2-deficient cells (Fig. 5A). Therefore, it was surprising to us that FXR2 associated with the SINV capsid in our proteomics screen, as well as a genome-wide screen for autophagy regulators^19^. Given that the FMR family of proteins are reported to be involved in autophagy and granulophagy ^26,27^ and autophagy is required for host defense against SINV, we examined whether FXR2 was required for clearing viral capsid. We found that FXR2-deficient cells accumulated more capsid. FXR2-deficient cells also showed reduced LC3-I-to-LC3-II conversion, fewer GFP-LC3 puncta, and persistent p62 abundance after SINV infection, consistent with impaired autophagy. This suggests that stress granule proteins may function as both proviral factors necessary for replication of alphavirus genomes and as antiviral factors that clear viral components via autophagy.

One possibility is that FXR2 couples recognition of the viral capsid to activation of the autophagy machinery. In this case, FXR2 either could act as a scaffold for capsid, p62, and LC3-associated proteins or by regulating the translation of factors required for selective autophagy. This interpretation is consistent with the identification of FXR2 in a genome-wide screen for autophagy regulators and with our observation that both FXR2 and p62 associate with immunoprecipitated capsid ^19^. Determining whether FXR2 binds capsid directly, whether this interaction depends on viral RNA, and which domains of FXR2 are required for capsid clearance will be important for defining its precise function in virophagy.

Through our PPI network analysis, we noted that FXR2 bridged two pathways: ribosomal large subunit biogenesis and core autophagy. The proteomics data suggests that FXR2 could potentially modulate autophagy initiation signals through its interactions with TRAF2 and PRKAA1 while simultaneously regulating ribosome production through its associations with RSL1D1 and RRP8. This dual functionality positions FXR2 as a potential regulatory node that could coordinate the balance between cellular protein synthesis and degradation pathways, which is critical for maintaining cellular homeostasis. In the context of viral infection, this bridging may regulate the synthesis of viral and host antiviral proteins, while also facilitating the degradation of viral proteins. Further work will be necessary to determine whether interactions with ribosomal proteins and/or autophagy proteins is necessary for the tissue protective role of FXR2 during SINV infection.

Several questions remain regarding the selectivity and generality of this pathway. Our experiments establish a requirement for FXR2 in autophagy induction, capsid clearance, cell survival, and protection from SINV encephalitis, but they do not yet distinguish whether capsid is the principal autophagic cargo responsible for the phenotype or whether FXR2 also regulates the turnover of additional viral or host components. It will therefore be important to confirm these findings with untagged viral proteins, primary neuronal cultures, and additional neurotropic alphaviruses, as well as to identify the cargo receptors and core autophagy proteins recruited by FXR2. Defining how capsid-associated FXR2 is partitioned between stress granules, ribonucleoprotein complexes, and autophagic structures may also explain how closely related RNA-binding proteins acquire opposing proviral and antiviral functions. Collectively, our results identify FXR2 as a previously unrecognized component of the protective autophagy response to alphavirus infection and suggest that enhancing selective viral-protein clearance could limit tissue injury even when viral replication itself is not reduced.

## Materials and Methods

### Cell culture

HeLa cells were obtained from the American Type Culture Collection (ATCC) and maintained in reduced-serum Opti-MEM I medium supplemented with 5% fetal bovine serum (FBS) and 1% antibiotic-antimycotic solution. BHK-21 cells used for plaque assays were maintained in Dulbecco’s Modified Eagle Medium supplemented with 10% FBS, 100 U/mL penicillin, and 100 µg/mL streptomycin

HeLa CRISPR FXR2 knockout (FXR2-KO) cell lines were generated using a CRISPR/Cas9-based genome editing approach following an established protocol ^28^. Briefly, the FXR2-KO cell line was generated in HeLa cells using a second-generation lentiviral vector system. The plasmids psPAX2 (Addgene #12260), pCMV-VSV-G (Addgene #8454), and lentiCRISPR v2 (Addgene #52961) were co-transfected into HEK293T cells using Lipofectamine 2000 (Thermo Fisher Scientific #11668027). The lentiCRISPR v2 plasmid carried a single-guide RNA (sgRNA; 5′-ACCTCCAGCTGACTATAATA-3′) designed to introduce a double-strand break within the FXR2 gene. Forty-eight hours after transfection, viral supernatants were collected, subjected to a freeze-thaw cycle for lysis, filtered through a 0.45-µm syringe filter to remove cellular debris. Following 48 hours of transduction, cells were selected using 2.0 µg/mL puromycin. Eight clones were selected for subsequent validation of FXR2 deficiency

To generate HeLa cells stably expressing GFP-LC3, cells were transfected with the pIRES.GFP-LC3.puro plasmid and selected with puromycin (1 µg/mL). FXR2 KO HeLa cells were reconstituted with wild-type FXR2 using a second-generation lentivirus system. The constructs psPAX2, p-CMV-VSV-g, and pLentip2a-BLAST (Addgene #211365, generously provided by the lab of Robert Orchard) were used to generate lentivirus, as above. The pLenti plasmid carried the FXR2 gene (Origene # RC209546, NCBI RefSeq. NM_004860.4) with a silent mutation to evade the sgRNA. FXR2 KO HeLa cells were transformed with this lentivirus and selected with 5.0 μg/mL blasticidin.

### Virus strains

SINV strain SVIA (ATCC) was derived from a low-passage isolate of the WT SINV strain AR339 ^29^. Recombinant SINV strain dsTE12Q engineered with a double subgenomic promoter was previously described was used for in vivo experiments for its neurovirulence ^30^. Recombinant SINV strain dsTE12Q.HA-capsid (SVIA.HA-capsid) and dsTE12Q.mCherry-capsid (SVIA.mCherry-capsid) was previously described ^1,6^. Viral stocks were propagated and titrated by plaque assays in BHK-21 cells.

### Mouse Strains

Fxr2 knockout mice were generated at Baylor College of Medicine from *Fxr2^tm^*^1a^ embryonic stem (ES) cells from the International Mouse Phenotyping Consortium (IMPC). The *Fxr2^tm^*^1a^ allele, on a C57BL/6N genetic background, contained LacZ and neomycin-resistance (Neo) selection cassettes inserted within intron 3 of Fxr2. ES cells were injected into blastocysts by the Genetically Engineered Rodent Models Core at Baylor College of Medicine to generate chimeric mice.

Chimeric mice were subsequently crossed with C57BL/6J mice to establish germline transmission of the targeted *Fxr2* allele. The resulting mice (*Fxr2^tm^*^2a^) were crossed with *ACTB:FLPe* transgenic mice (The Jackson Laboratory, Stock No. 005703) to mediate FLP recombinase-dependent excision of the FRT-flanked LacZ and Neo cassettes. This breeding generated the *Fxr2^tm^*^2c^ allele, in which the LacZ and Neo cassettes were removed while loxP sites flanking *Fxr2* exons 4 and 5 were retained.

To generate the *Fxr2^tm^*^2d^ knockout allele, *Fxr2^tm^*^2c^ mice were crossed with *EIIa-Cre* transgenic mice (The Jackson Laboratory, Stock No. 003725). Cre-mediated recombination resulted in deletion of *Fxr2* exons 4 and 5, and germline transmission of the recombined *Fxr2^tm^*^2d^ allele was established. *Fxr2^tm^*^2d^ mice were used for all subsequent mouse experiments.

*Fxr2^Tm^*^2d^ (“FXR2 KO”) mice were maintained at UT Southwestern by breeding heterozygotes to generate mixed-genotype litters for animal experiments. Animals were raised and manipulated in barrier facilities. All animal work was registered under an animal protocol approved by the UT Southwestern Institutional Animal Care and Use Committee.

### Western blot analysis

Cells were lysed in RIPA buffer (Cell Signaling Technology #9806) supplemented with protease and phosphatase inhibitor cocktails. Protein samples were separated on 4-20% gradient SDS-polyacrylamide gels (Bio-Rad) and transferred onto PVDF membranes. Membranes were blocked with 5% nonfat dry milk in TBS containing 0.05% Tween-20 (TBST) and incubated with the primary and secondary antibodies listed below. Protein bands were visualized using enhanced chemiluminescence (Thermo Fisher Scientific) and imaged using a Bio-Rad ChemiDoc system.

| Protein | Vendor | Catalog Number | Dilution |
| --- | --- | --- | --- |
| HA-tag | Cell Signaling Technologies | 3724 | 1:1000 |
| SINV Capsid | Gift from Diane Griffin | -- | 1:5000 |
| HA-tag | Sigma-Aldrich | 11867423001 |  |
| P62 | Abnova | 8878 | 1:2000 |
| Actin-HRP | Santa Cruz Biotechnology | sc-47778 HRP | 1:10,000 |
| LC3II | Novus | 2220 | 1:1000 |
| FXR2 (human) | Santa Cruz Biotechnology | sc-166433 | 1:1000 |
| Vinculin | Cell Signaling Technologies | 13901 | 1:1000 |
| Lamin B1 | Cell Signaling Technologies | 13435 | 1:1000 |
| FXR2 (mouse) | Thermo Fisher Scientific | MA1-5773 |  |
| GAPDH | Thermo Fisher Scientific | MA5-15738 |  |

### Co-immunoprecipitation and proteomic profiling to identify capsid-interacting host proteins

To systematically identify host proteins that interact with the capsid protein of Sindbis virus (SINV) Infection. 1×10^7^ HeLa cells were infected with SVIA.HA-capsid MOI=10 for 1h and 7h, followed by immunoprecipitation with anti-HA antibody. After protein isolation, protein lysates were pre-cleared with protein G PLUS agarose beads (SCBT, sc-2002) for 1.5h at 4℃ with gentle agitation. After pelleting the beads, supernatants were mixed with 0.8 µg/mL of rat anti-HA antibody (Sigma-Aldrich, 11867423001) and 40μl of protein G PLUS agarose beads and then incubated at 4℃ overnight with gentle agitation. Agarose beads were pelleted by centrifugation at 3000g for 1 minute followed by 3 washes with ice-cold lysis buffer. Immunoprecipitated proteins were eluted by adding 2X Laemmli sample buffer (Bio-Rad Laboratories) containing 5% β-mercaptoethanol (Bio-Rad Laboratories) and boiling sample for 5 minutes followed by coomassie staining and western blot with anti-HA antibody. These samples were subjected to LC-MS/MS for proteomics analysis performed by UT Southwestern Proteomics Core. Raw MS data files are analyzed using Proteome Discoverer v2.4 SP1 (Thermo), with peptide identification performed using Sequest HT searching against the human protein database from UniProt. Fragment and precursor tolerances of 10 ppm and 0.6 Da are specified, and three missed cleavages were allowed.

### Analysis of LC/MS proteomics data

#### Data Integration and Preprocessing

Proteomics data from Batch A (1 experiment with control, 1 HPI and 7 HPI samples) and Batch B (2 experiments, each with control, 1 HPI and 7 HPI samples) were merged and underwent log2 transformation followed by quantile normalization using the normalize BetweenArrays function from the limma package to reduce technical variation. To mitigate systematic differences between batches while preserving biological variation, we implemented a mean-centering batch correction approach. For each protein detected across multiple batches, correction factors were calculated for each experimental condition separately. Using Batch A as the reference, we computed the mean difference between batches for each condition and applied this correction to adjust Batch B values accordingly. This condition-specific correction preserves within-condition relationships while reducing between-batch systematic differences, as visualized through Principal Component Analysis before and after correction.

Following our relaxed intersection approach, we retained proteins with at least one observation per experimental condition. For proteins with limited missingness (≤2 samples), we performed condition-specific mean imputation, where missing values were replaced with the mean abundance of the protein within the same experimental condition. We specifically employed this conservative condition-specific mean imputation approach because the high missingness in Batch A made neighborhood-based methods like KNN unreliable, and the absence of replicates in Batch A prevented variance estimation required for model-based imputation.

#### Differential Expression Analysis

Differential expression analysis was performed using the limma package by fitting a linear model to the normalized, batch-corrected, and selectively imputed data. The design matrix incorporated both condition (Control vs. Infection) and time point (1h and 7h) information. Empirical Bayes moderation of standard errors was applied to improve statistical robustness, particularly important for datasets with limited replication. Proteins with adjusted p-values < 0.05 were considered significantly differentially expressed.

#### Gene Ontology Enrichment Analysis

For proteins overlapping with the Orvedahl 2010 pape^19^, Gene Ontology (GO) enrichment analysis was conducted using the clusterProfiler package. Significantly differentially expressed proteins were mapped to gene symbols using a combination of annotations from the original dataset and extraction from protein descriptions where necessary. ENTREZ IDs were obtained using the mapIds function from the org.Hs.eg.db package. GO enrichment was performed using the enrichGO function with the Biological Process (BP) ontology, with the entire detected proteome serving as the background reference set. Significant GO terms were identified using a p-value cutoff of 0.05 and Benjamini-Hochberg correction for multiple testing.

#### FXR2 Time Course Analysis

To analyze FXR2 protein expression dynamics during infection, we normalized FXR2 abundance relative to a panel of stable reference proteins. Three established housekeeping proteins (YWHAZ, PPIA, and GAPDH) were selected as internal controls for normalization. For each sample, we calculated the geometric mean of these three housekeeping proteins’ normalized abundance values.

FXR2 relative abundance was determined by dividing its normalized values by the housekeeping protein geometric mean for each corresponding sample. To facilitate interpretation of expression changes over time, fold changes were calculated relative to the mean expression in control samples (t=0h). Statistical significance of temporal changes in FXR2 expression was assessed using linear modeling with the limma package. The design matrix incorporated both time points (0h, 1h, and 7h) and batch information. One-sided hypothesis testing was performed to evaluate increases in FXR2 expression at 1h and 7h post-infection relative to control (0h) samples.

The temporal dynamics of FXR2 expression were visualized in a time course plot showing fold changes at 0, 1, and 7 hours post-infection. Individual data points were plotted with color-coding by condition, mean values connected by a trend line, standard error bars to indicate variability, and a reference line at fold change = 1 representing baseline expression. Statistical significance was indicated with horizontal bars and asterisks above the relevant time point comparisons.

#### Protein-Protein Interaction (PPI) Network Plot

Human protein–protein interactions were obtained from BioGRID (version 4.4.238). We first defined a node set comprising all significantly differentially expressed proteins (adj. p-value < 0.05, |fold change| > 2.0), core autophagy and autophagy-associated proteins as defined by Orvedahl et al.^19^, and our protein of interest, FXR2. We retained every BioGRID interaction in which both partners belonged to this node set, regardless of the order in which they were listed (A–B or B–A). Interactions were then treated as undirected, and duplicate edges and self-loops were removed to produce the final network.

Nodes were assigned to one of four groups: core autophagy, autophagy-associated, significantly upregulated, or protein of interest. Downregulated proteins were excluded from the visualization to improve interpretability. Significantly upregulated proteins not assigned to any other group were further annotated with k-means clustering (k = 3) in STRING (https://string-db.org/). The network was exported in JSON format and visualized with a HTML/JavaScript application built on Cytoscape.js.

### Autophagy assay

To assess autophagy induction in HeLa cells, the number of autophagosomes, represented by GFP-LC3 puncta per cell, was quantified by fluorescence microscopy by an observer blinded to both treatment condition and cell type. Autophagy was further evaluated by measuring LC3-I to LC3-II conversion by immunoblotting using an anti-LC3B antibody. For virophagy experiments, cells were infected with Sindbis virus (SINV) strains at a multiplicity of infection (MOI) of 10 in Opti-MEM I Reduced Serum Medium supplemented with 5% FBS for 1 h at 37°C. Following infection, the inoculum was removed and replaced with fresh culture medium, while mock-infected cells were maintained in culture medium alone. Autophagic flux was assessed by treatment with 100 nM bafilomycin A1 (BafA1) for 2 h.

### SINV growth curves

To evaluate viral replication kinetics using a multistep growth curve assay, HeLa cells were seeded in triplicate in 6-well plates and infected at a multiplicity of infection (MOI) of 0.01 for 1.5 h. Following adsorption, the inoculum was removed and replaced with fresh culture medium. At the indicated time points, 100μL of supernatant was collected from each well, and viral titers were quantified by plaque assay on BHK-21 cells.

### Cell death assay

To assess virus-induced cell death, cells were either mock infected or infected with SINV MOI=10 for 24 h. Cell viability was subsequently measured using the CellTiter-Glo Luminescent Cell Viability Assay (Promega, G7570) according to the manufacturer’s protocol. Luminescence was recorded using a CLARIOstar microplate reader (BMG Labtech), and viability was determined relative to mock-infected controls.

### Survival and titer of mice infected with SVIA

Seven-day-old *Fxr2^-/-^*, *Fxr2^+/-^*, and *Fxr2^+/+^* littermates were infected with 1 × 10³ PFU of the dsTE12Q strain in a 30 μL inoculum^30^. Mice were inoculated intracerebrally in the right cerebral hemisphere with SVIA diluted in Hank’s Balanced Salt Solution (HBSS; Thermo Fisher Scientific). For survival studies, animals were monitored daily for 21 days following infection. For virological and histological analyses, animals from each genotype were randomly selected for tissue collection at 1, 4, and 7 days post-infection. The right cerebral hemisphere was rapidly frozen in liquid nitrogen, homogenized in HBSS, and analyzed by plaque assay to quantify viral titers. The contralateral hemisphere was fixed in 4% paraformaldehyde, cryoprotected in 30% sucrose, and embedded in paraffin for subsequent histological analysis.

### Histology

The UT Southwestern Histology Core embedded the fixed tissue into paraffin, took Paxinos 110 sections, and stained sections with hematoxylin and eosin (H&E). A UT Southwestern veterinarian pathologist, Dr. Mary Carter, scored blinded H&E slides, rating inflammation and cell degeneration and/or apoptosis in the hippocampus and cortex.

For in vivo assessment of cell death, TUNEL staining was performed on paraffin-embedded sagittal brain sections using the ApopTag Peroxidase In Situ Apoptosis Detection Kit (EMD Millipore, S7100) according to the manufacturer’s instructions. Whole-brain sections were imaged using a Zeiss Axio Scan.Z1 slide scanner with a PLAN-APOCHROMAT 20×/0.8 NA objective (Carl Zeiss Microscopy; UT Southwestern Whole Brain Core Facility). TUNEL slides were scanned and quantitated using the Aperio ImageScope software.

## Statistics

Statistical analyses were performed using GraphPad Prism version 10. Statistical tests are indicated in the corresponding figure legends. For in vitro experiments, comparisons between two groups were performed using an unpaired, two-tailed Student’s *t*-test, while comparisons among more than two groups were performed using analysis of variance (ANOVA). For in vivo pathogenesis studies, differences between two groups were assessed using the nonparametric Mann–Whitney *U* test. Survival data were analyzed by generating Kaplan–Meier survival curves and comparing groups using the log-rank test. Image quantification was performed using AxioVision Rel. 4.8 software with a pre-defined analysis macro. Cell counting was performed by an observer blinded to experimental condition and genotype. Immunofluorescence microscopy (IFM) analysis was conducted using ImageJ software.

## Data availability

The mass spectrometry proteomics data have been deposited to the ProteomeXchange Consortium via the MassIVE repository with the dataset identifier PXD086131 and MassIVE identifier MSV000103691.

## Author Contributions

VS, JWT, TAR conceptualized the study. VS, PD, JWT, TAR analyzed the data. TAR acquired funding. VS, PD, ZZ, NC performed experiments. YG and DLN made *Fxr2^-/-^* mice. WC and TW provided bioinformatic analysis. TAR provided project administration. TAR, VS generated figures. TAR wrote the original draft. The manuscript was edited by all authors.

## Acknowledgements

The UT Southwestern Histology Core performed embedded tissue and stained sections. The UT Southwestern Proteomics Core performed proteomic analysis. We thank Dr. Mary Carter of UT Southwestern for histology scoring. We thank Diane Griffin for the anti-capsid antibody, and Stacey Crockett and Robert Orchard for reagents. This work was supported by NIH grants U19 AI142784 (TAR), 2R01 AI130020-08 (TAR) and Welch grant I-2157-20230405 (TAR). Tiffany Reese holds the Paul and Betty Meek – FINA Professorship in Molecular Immunology at UT Southwestern.

